# Hyperhomocysteinemia does not increase the risk of intracerebral hemorrhage in hypertensive mice

**DOI:** 10.64898/2026.01.28.702446

**Authors:** Hui Zhao, Ning Hou, Xiao Shi, Zhaoyun Liu, Shouying Ding, Tonghui Wang, Qiang Feng

## Abstract

**Objective:** Hyperhomocysteinemia (HHcy) affects approximately 75% of the population in China, and there is currently controversy regarding whether HHcy increases the risk of hemorrhagic stroke. This study aims to investigate the effects of high homocysteine (Hcy) levels on cerebral hemorrhage in hypertensive mice by administering homocysteine to them.

**Methods:** Male C57BL/6 mice at 8 months of age were used in the experiment. The study was divided into two groups: the Hcy + AngII + L - NAME group and the AngII + L - NAME group. Magnetic resonance imaging (MRI) was performed when the mice exhibited signs of cerebral hemorrhage.After the hemorrhage, anesthesia was induced to euthanize the animals, and then the brain tissue was fixed. The total rearing period was 18 weeks. The relationship between homocysteine and stroke was described by plotting survival curves. The location and quantity of cerebral hemorrhage were determined through histopathological staining.

**Results:** The serum Hcy concentration of mice fed with Hcy for 6 weeks increased to 23.07 μmol/L, and the blood pressure ranged from 170 to 180 mmHg. The number of deaths due to cerebral hemorrhage was 10 in both the AngII + L - NAME + Hcy group and the AngII + L - NAME group. The p - value of the survival curves between the two groups was 0.162, indicating no statistically significant difference.

**Conclusion:** The results demonstrated that elevated homocysteine levels did not influence the incidence of intracerebral hemorrhage in hypertensive mice.

Hyperhomocysteinemia does not increase the risk of intracerebral hemorrhage in hypertensive mice

---

Hypertension is the most significant risk factor for intracerebral hemorrhage (ICH) [1]. Among adult hypertensive patients in China, approximately 75% also exhibit HHcy (Hcy ≥10 μmol/L), a condition known as HHcy[2]. Epidemiological studies have demonstrated that patients with type HHcy have a significantly increased risk of stroke, particularly in ischemic stroke research[3-6]. However, studies on the relationship between HHcy and ICH are relatively limited, and the findings remain controversial. Some studies suggest that HHcy is an independent risk factor for ICH. A cohort study from northern India demonstrated that homocysteine deficiency is associated with hemorrhagic stroke [7]. Studies suggest that Hcy levels ≥ 10 μ mol/L are risk factors for recurrent hypertensive ICH [8]. Research has demonstrated a positive correlation between Hcy levels and the severity of cerebral microbleeds [9,10]. Plasma total homocysteine (tHcy) is an independent influencing factor for cerebral microbleeds, and it shows a positive correlation with the number of hemorrhage sites [11]. A study using inverse variance weighting (IVW) analysis found that genetically elevated homocysteine levels were associated with an increased risk of ICH; however, the final results were not statistically significant [12]. However, some studies indicate that HHcy has no impact on the occurrence of hemorrhagic stroke [13-14]. A cohort study found that high levels of Hcy (≥11.0 μmol/L) were not significantly associated with an increased risk of hemorrhagic stroke compared to low levels (<7.0 μ mol/L)[15]. A meta - analysis by Yusheng He et al. found no association between blood homocysteine (Hcy) levels and the risk of cerebral hemorrhage[16].

This uncertainty casts doubt on the efficacy of interventions targeting hyperhomocysteinemia, such as folic acid supplementation, in preventing ICH. Therefore, a mouse model of hypertension was established[17, 18]. Concurrently, mice were fed a diet to induce hyperhomocysteinemia (HHcy) to investigate its impact on hemorrhagic stroke in hypertensive mice..

## Methods

### 1. Construction of Hypertensive Mouse Models

Eight - month - old male C57BL/6 mice (Beijing VTRILIA Laboratory Animal Co., Ltd.) were divided into two groups: the Hcy + AngII + L - NAME group and the AngII + L - NAME group, each consisting of 20 mice. The Hcy + AngII + L - NAME group was provided with water containing 1.8 g/L homocysteine. At week 7, an ALZET micro - osmotic pump containing angiotensin II (AngII, 1,000 ng/kg/min) was implanted into the subcutaneous tissue of the mice’s backs, and they were also administered 100 mg/kg/day of L - NAME. The AngII + L - NAME group was not provided with water containing Hcy. The tail blood pressure of the mice was measured using a BP - 2000.

All experiments were reviewed and approved by the Laboratory Animal Committee of Tai’an Municipal Hospital (Ethical Approval Number: 20210328-6).

### 2. Measurement of Homocysteine Concentration in Mouse Blood

After 6 weeks of water feeding with 1.8 g/L homocysteine, the mice were anesthetized, blood was collected via cardiac puncture, and plasma Hcy concentration was detected by an automatic biochemical analyzer.

### 3. Assessment of ICH

According to the “Behavioral Assessment Methods for Mice with ICH”, behavioral assessments were conducted three times a day (morning, noon, and evening) on the mice. Behavioral signs of ICH included contralateral forelimb extension, circling behavior, tremors, paralysis, or other motor dysfunctions. When ICH behavior was observed, the mice were anesthetized with isoflurane and placed on an MRI machine for brain flat-plate scanning (resolution: 1 mm) to determine the hemorrhage site and volume. The mice were then euthanized under anesthesia, and blood and brain tissue were collected.

### 4. Detection of the number of bleeding sites and the area of bleeding

The mouse brain was sectioned at a thickness of 3μm, and hemorrhagic sites and areas were identified by hematoxylin and eosin (H&E) staining at intervals of 10 sections. Photographs were taken of the hemorrhagic sites, and the size of the hemorrhagic areas in the mice was analyzed using the Image-Pro Plus (IPP) image processing and analysis software.

### 5. Detection of vascular smooth muscle quantity at the bleeding site

Adjacent slices of the hemorrhage were selected for immunofluorescence staining of vascular smooth muscle.After taking photos using a laser confocal microscope, the number of smooth muscle cells was analyzed.

### 6. Data Analysis

All measurement data were expressed as mean ± standard deviation. Chi-square tests and non-parametric tests were employed to analyze differences between groups. A p - value < 0.05 was regarded as statistically significant. The cumulative incidence and survival rates of cerebral hemorrhage were analyzed using the K - M survival curve. Data were analyzed using SPSS 19.0.

## Results

### 1. Basic Information of Mice

After 6 weeks of feeding with 1.8 g/L homocysteine, the average serum homocysteine concentration was 23.07 μmol/L. One week after subcutaneous embedding of angiotensin II in the dorsal skin of mice, the blood pressure rapidly increased to 160 mmHg and eventually stabilized between 170 and 180 mmHg. No significant difference was observed between the two groups (Figure A).

**Figure A:**
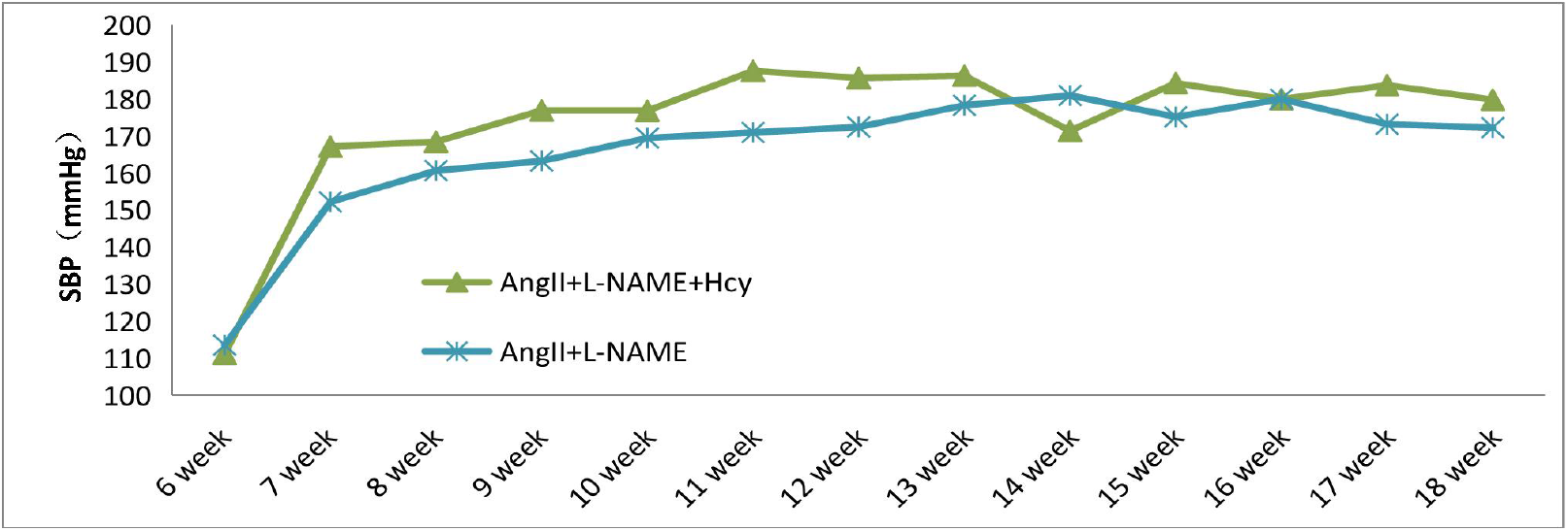
Mean systolic blood pressure (SBP) within the group.

Behavioral assessments, MRI imaging (Figures B, C), and pathological examinations revealed that most mice died from ICH (Figures D, E), with the white hyperintense areas indicating hemorrhagic sites. Figure B demonstrates MRI-detected hemorrhages in the basal ganglia, while Figure C shows cortical hemorrhages. Figures D and E reveal diffuse hemorrhages from the vasculature after hematoxylin - eosin (HE) staining. These findings confirm the success of our ICH model.

**Figure B:**
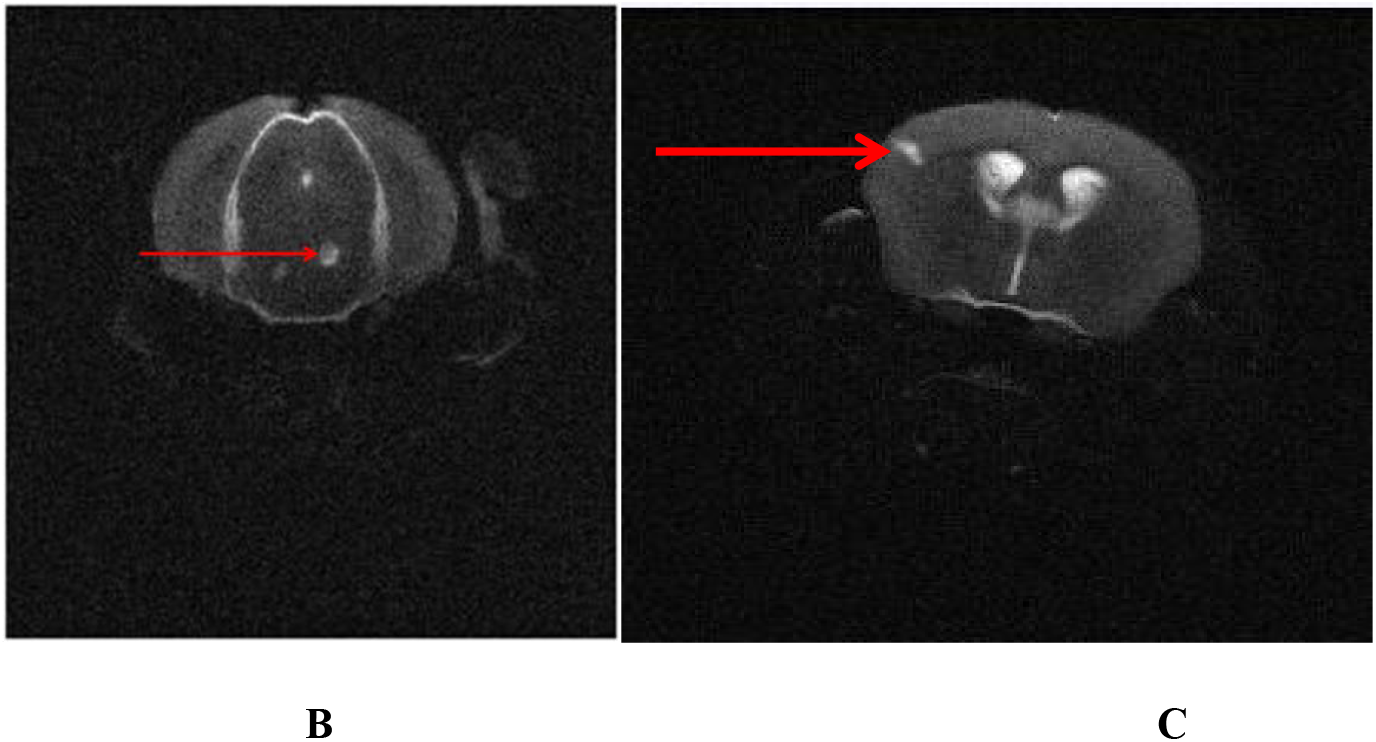
MRI shows hemorrhagic lesions near the brainstem, with the hemorrhagic focus indicated by the arrow. Figure C: MRI detection reveals hemorrhagic lesions near the cerebral cortex, with the arrow indicating the hemorrhagic focus.

**Figures D and E:**
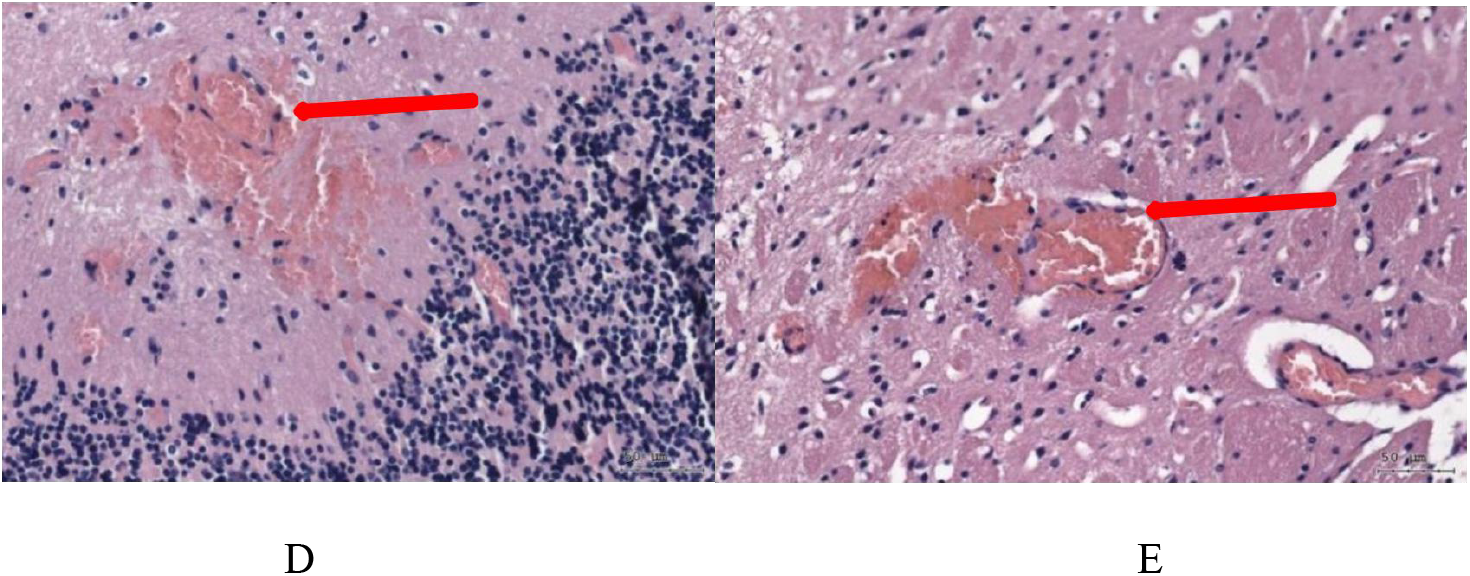
Diffuse hemorrhagic foci near the cerebral vessels were identified by H&E staining. At the arrow location, the hemorrhagic vessel is visible, and blood is flowing out of it.

### 2. Mortality and survival curves of mice

After 18 weeks of feeding, all mice were euthanized, and the causes of death in each group were recorded. In the AngII + Hcy + L - NAME group, 10 mice died from ICH, 1 from peritoneal hemorrhage, and 3 due to perioperative mortality. In the AngII + L-NAME group, 10 mice died of ICH, 1 from thoracic hemorrhage, 1 from peritoneal hemorrhage, and 3 from perioperative mortality (Table 1).

**Table 1:**
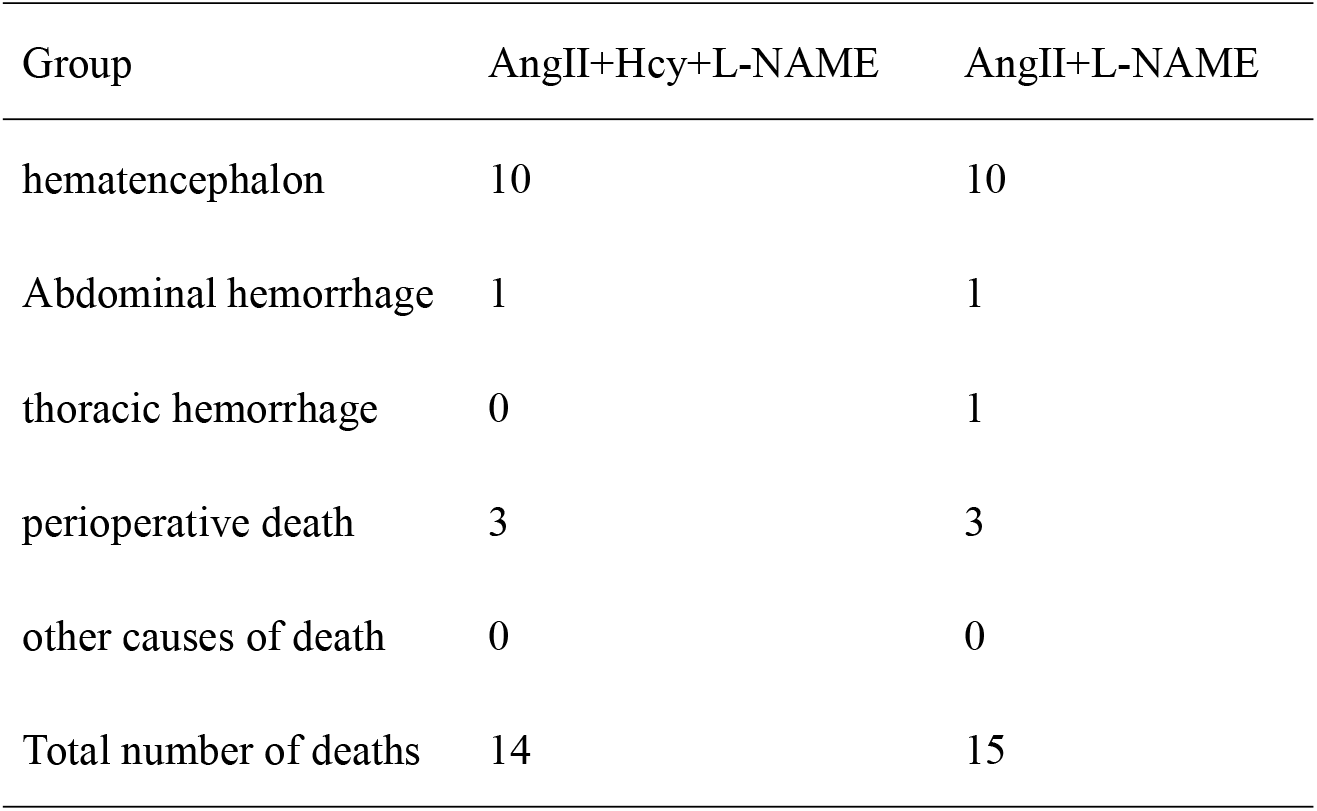
Mortality and types of death in mice.

| Group | AngII+Hcy+L-NAME | AngII+L-NAME |
| --- | --- | --- |
| hematencephalon | 10 | 10 |
| Abdominal hemorrhage | 1 | 1 |
| thoracic hemorrhage | 0 | 1 |
| perioperative death | 3 | 3 |
| other causes of death | 0 | 0 |
| Total number of deaths | 14 | 15 |

After excluding deaths from other causes and retaining only mice that died from ICH, the survival curves showed no inter - group differences between the Hcy + AngII group and the AngII group (*p* = 0.162). Similarly, no inter-group differences were observed between the Hcy + AngII + L–NAME group and the AngII + L–NAME group (*p* = 0.918) (Figure H).

**Figures h:**
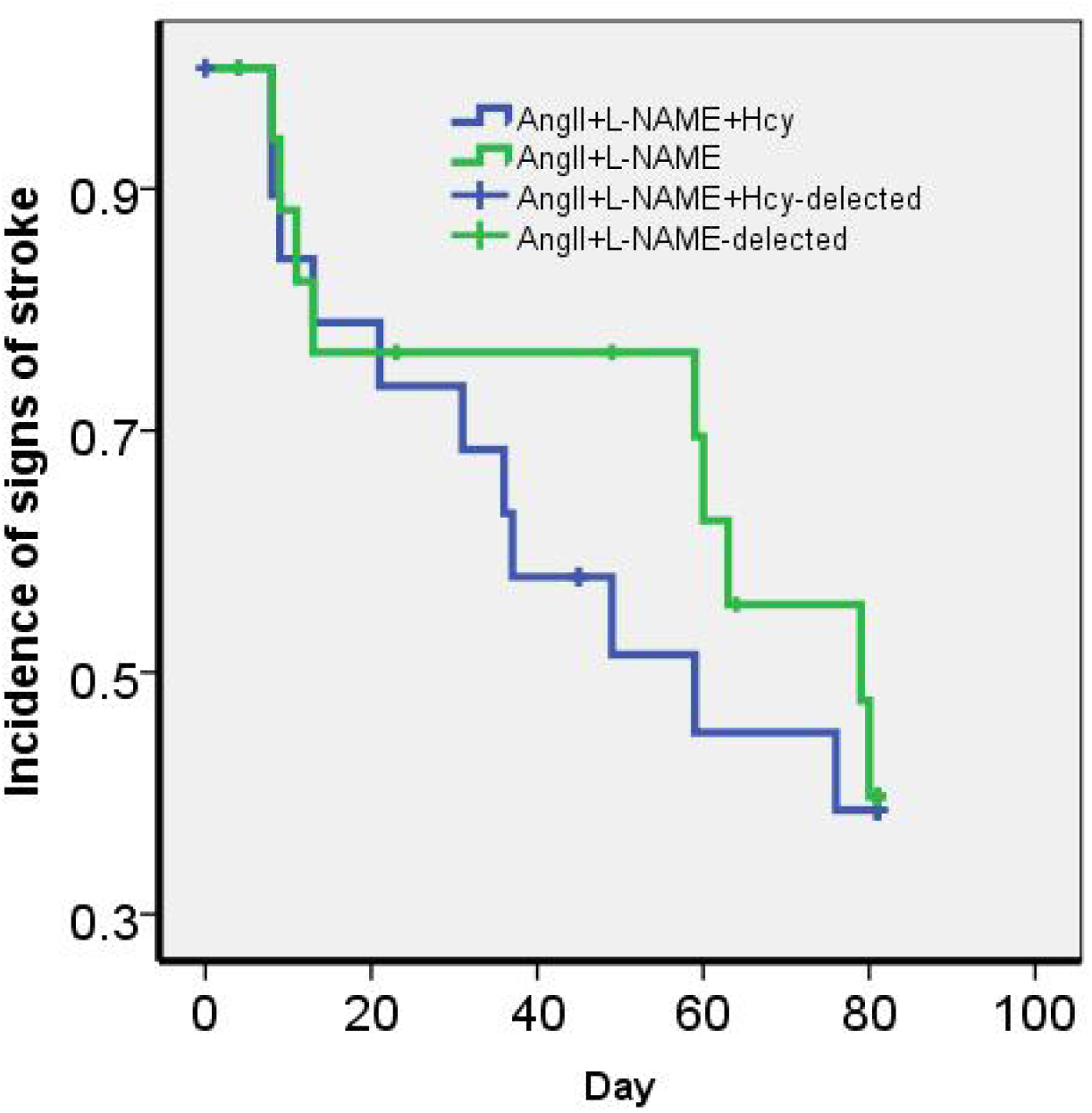
K- M curve of death from ICH in mice: Starting from day 0 when AngII was subcutaneously implanted.

### 3. Number of petechiae and bleeding area

Three brains from each group of mice with cerebral hemorrhage were randomly selected for section analysis. After HE staining, the average number of hemorrhagic sites per group was found to be as follows: 10.7 in the Hcy + AngII + L - NAME group and 10.3 in the AngII + L - NAME group. A comparison of the number of hemorrhagic sites between the Hcy + AngII + L - NAME group and the AngII + L - NAME group revealed a p - value of 0.64, indicating no statistically significant difference.

After measuring the hemorrhagic area at the bleeding site using the IPP software, the hemorrhagic areas in each group were as follows: Hcy + AngII group: 33,227.37 μm^2^; Hcy + AngII + L - NAME group: 25,884.69 μm^2^. The difference in hemorrhagic area between the Hcy + AngII + L - NAME group and the AngII + L - NAME group was statistically significant (*p* = 0.003) (Table 2).

**Table 2:** Number of bleeding sites and size of bleeding area.

| Group | AngII+Hcy+L-NAME | AngII+L-NAME |
| --- | --- | --- |
| Number of mice | 3 | 3 |
| Number of bleeding sites | 32 | 31 |
| Number of bleeding sites per mouse<br>on average | 10.7 | 10.3 |
| P price | 0.64 |  |
| Average bleeding area ($\mu\text{m}^2$) at all<br>bleeding sites | 2426.68 | 6040.92 |
| Average hemorrhage area per mouse<br>($\mu\text{m}^2$) | 25884.69 | 60409.30 |

### 4. Number of vascular smooth muscle cells at the bleeding site

After immunofluorescence staining and software analysis, the average number of smooth muscle cells in each group was calculated as follows: AngII + L–NAME + Hcy group: 7.92; AngII + L – NAME group: 8.15. No significant difference was observed between the Hcy + AngII + L–NAME group and the AngII + L–NAME group (*p* = 0.451; Figure I).

**Figure I:**
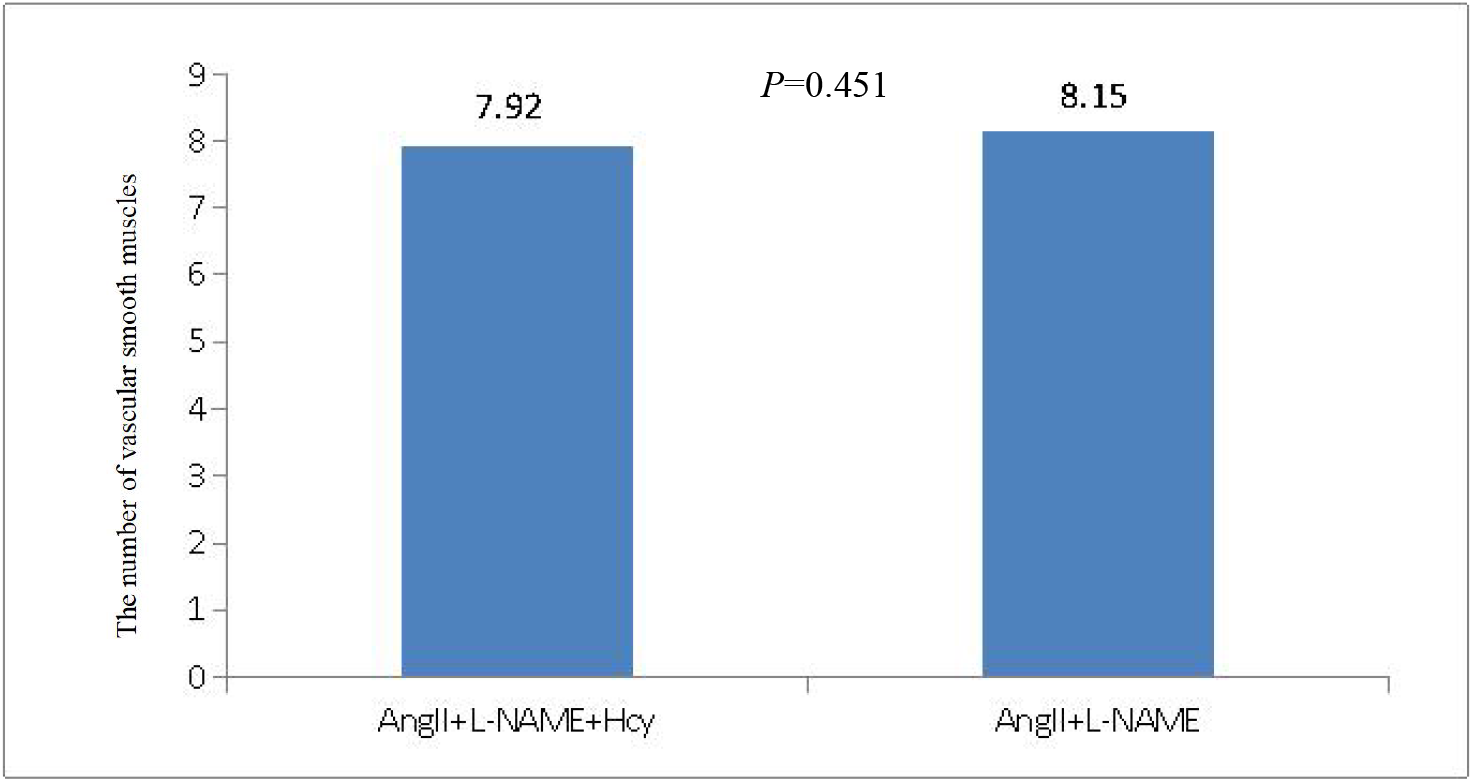
Average number of vascular smooth muscle cells per vessel

## Discussion

This study is the first experimental investigation at the animal level, both in China and abroad, to examine the relationship between homocysteine and hypertensive ICH. The findings revealed that elevated homocysteine levels did not influence the occurrence of ICH in hypertensive mice.

Numerous studies have demonstrated that homocysteine can damage vascular endothelial cells, induce structural changes in blood vessels, and disrupt their normal functions[19-22]. Homocysteine undergoes in vivo metabolism to produce hydrogen sulfide, a potent vasodilator and antioxidant[23]. The metabolites of homocysteine in vascular endothelial cells can affect smooth muscle cells, leading to vascular dysfunction and subsequently causing hypertension[24]. Studies have shown that HHcy is associated with thrombus formation, as it can affect the binding of thrombin, protein C, and thrombin regulatory proteins, thereby leading to thrombus formation.[25]. Homocysteine can promote the binding of lipoprotein A (LpA) to fibrin and fibrinolytic proteins, thereby enhancing the atherogenic potential of LpA[26-28]. Meanwhile, homocysteine plays a significant role in promoting the proliferation of vascular smooth muscle cells[29,30]. Homocysteine promotes the proliferation of smooth muscle cells by binding to receptors associated with homocysteine redox reactions in smooth muscle cells, thereby influencing the redox reactions of smooth muscle cells and their surrounding cells[31,32]. Studies have also demonstrated that Hcy increases ADP levels in endothelial cells by inhibiting ADPase activity. The elevated ADP levels enhance platelet activity, which subsequently promotes thrombus formation[33]. The aforementioned mechanisms are primarily associated with thrombosis and atherosclerosis. However, the occurrence of ICH is predominantly attributed to vascular thinning and the disappearance of smooth muscle[34,35]. These findings are contrary to the pathogenesis of ICH. From this perspective, the explanation of Hcy’s potential involvement in ICH remains inconclusive.

The relationship between the number of vascular smooth muscle cells and vascular tone is direct[36,37]. This study found that the number of vascular smooth muscle cells in the cerebral hemorrhage group was reduced compared with that in the hypertension group, but no statistically significant difference was observed. This suggests that homocysteine (Hcy) does not influence the occurrence of cerebral hemorrhage in mice by reducing the number of smooth muscle cells. This finding is consistent with our survival curve results. We need to identify more hemorrhagic sites to study their smooth muscle cell counts and demonstrate the relationship between smooth muscle and cerebral hemorrhage.

The core finding of this study, namely that HHcy does not exacerbate the risk of ICH in hypertensive mice, is consistent with some clinical observations. This result suggests that the role of HHcy in the pathophysiological process of ICH may be background-dependent. In the context of hypertension, a dominant risk factor that directly causes structural damage to cerebral blood vessels (e.g., reduction of vascular smooth muscle cells and disruption of the intima - media elastic lamina), the molecular mechanisms mediated by HHcy — mainly associated with atherosclerosis and thrombosis (e.g., endothelial dysfunction, oxidative stress, and procoagulant states)—may not be the key drivers of ultimate vascular rupture. This may explain why critical indicators such as survival curves, the number of hemorrhagic sites, and the count of vascular smooth muscle cells at the hemorrhagic sites showed no statistically significant differences between groups.

This study has certain limitations. The small sample size in each group may introduce statistical errors. Future research should employ larger populations and conduct more detailed investigations to fully elucidate the relationship between HHcy and hypertensive ICH. Additionally, the L - NAME - induced ICH model used in this study has inherent limitations, and its consistency with other ICH models requires further investigation.

## Nonstandard Abbreviations and Acronyms

HHcy: Hyperhomocysteinemia
Hcy: homocysteine
MRI: Magnetic Resonance Imaging
ICH: intracerebral hemorrhage
IVW: nverse variance weighting
H&E: hematoxylin and eosin
IPP: Image-Pro Plus
LpA: lipoprotein A

## Acknowledgments

We express our gratitude to the authors of this article for their contributions to this paper, as well as to the funding providers of this study.

